# Neopolyploidy-induced changes in the giant duckweed (*Spirodela polyrhiza*) alter herbivore preference, performance, and plant population performance

**DOI:** 10.1101/2023.11.14.567047

**Authors:** Hannah R. Assour, Tia-Lynn Ashman, Martin M. Turcotte

**Author notes:** **Corresponding author**: Hannah Assour.

## Abstract

**Premise:** Polyploidy is a widespread mutational process in angiosperms that may alter population performance of not only plants but also their animal associates. Yet, knowledge of whether ploidy affects plant-herbivore dynamics is scarce. Here, we test whether aphid herbivores exhibit preference for diploid or neopolyploid plants, whether ploidy impacts plant and herbivore performance, and whether these interactions depend on plant genetic background.

**Methods:** Using multiple pairs of independently synthesized neotetraploid greater duckweed (*Spirodela polyrhiza*) and their diploid progenitors, we evaluated the effect of neopolyploidy on duckweed’s interaction with the water-lily aphid (*Rhopalosiphum nymphaeae*). Using two-way choice experiments, we first evaluated feeding preference by the herbivore. We then evaluated the consequences of ploidy on aphid and plant performance by measuring population growth over multiple generations.

**Key Results:** Aphids preferred neopolyploids over diploids when the plants were provided at equal abundances but not when they were provided at equal surface area, indicating the role of plant size in driving this preference. Additionally, neopolyploidy increased aphid population performance, but this result was highly dependent on the genetic lineage of the plant. Lastly, the impact of herbivory on neopolyploids vs. diploid duckweed varied greatly with genetic lineage, but overall, neopolyploids appeared to be generally less tolerant than diploids.

**Conclusions:** We conclude that polyploidization can impact the preference and performance of herbivores on their plant hosts, whereas plant performance depends on complex interactions between herbivory, ploidy, and genetic lineage. These results have significant implications for the establishment and persistence of plants and herbivores in nature.

## INTRODUCTION

Plant polyploidy, or whole genome duplication, is a dramatic and prevalent mechanism of differentiation in plants (Nuismer and Thompson, 2001; Thompson et al., 2004; Arvanitis et al., 2010; Ramsey and Ramsey, 2014; Segraves and Anneberg, 2016). Thirty-five percent of extant angiosperms are of recent polyploid origin, and all angiosperms have at least one whole-genome duplication event in their evolutionary past (Soltis et al., 2009; Wood et al., 2009; Jiao et al., 2011). Polyploidy is extremely widespread; polyploids are found across the globe, with some terrestrial biomes containing as high as 51% polyploid frequency and mixed-ploidy species (exhibiting both diploid and polyploid cytotypes) are also common (Kolář et al., 2017; Rice et al., 2019). Similarly, 52% of aquatic plant species are thought to be either polyploid or mixed-ploidy species (Magalhães et al., 2021). Researchers have long been trying to explain this apparent enhanced ecological and evolutionary success of polyploids, particularly by investigating the genotypic and phenotypic differences of polyploids from their diploid progenitors (Comai, 2005; Madlung, 2013; Levin and Soltis, 2018; Fox et al., 2020; Van-de-Peer et al., 2020). However, while studies of the mechanisms behind polyploid success in terrestrial species have been accumulating over the last few decades, comparatively less is known for aquatic species.

Polyploid plants often differ from their diploid ancestors in a variety of ways that can impact their interaction with abiotic and biotic factors (Gross and Schiestl, 2015; Wei et al., 2019; Clo and Kolář, 2021). It is argued polyploidy enhances tolerance to abiotic stressors, such as heat, cold, salt, and nutritional stress (Yang et al., 2014; Godfree et al., 2017; Song et al., 2020; Tossi et al., 2022; Anneberg et al., 2023a). Yet, tolerance to biotic stressors has received much less attention, despite the likelihood that phenotypic differences also impact species interactions (Segraves and Anneberg, 2016; Forrester and Ashman, 2017; Rezende et al., 2020; Anneberg et al., 2023b). Specifically, polyploidy-induced phenotypic and genotypic changes can lead to novel interactions with other species, such as herbivores, pollinators, and microbes (Arvanitis et al., 2010; McCarthy et al., 2016; Porturas et al., 2019; Walczyk and Hersch-Green, 2019; Forrester et al., 2020; Rezende et al., 2020; Curé et al., 2022; Anneberg et al., 2023b).

Numerous changes due to polyploidy including plant body size, trichome size/number, leaf thickness, secondary metabolite production, and cellulose content may alter herbivore performance and preference as well as plant tolerance and resistance to herbivores (Bagheri and Mansouri, 2015; Corneillie et al., 2018; Wei et al., 2019; Bomblies, 2020; Hamarashid et al., 2022; Malacrinò et al., 2022). For example, the gigas effect, or the enlargement of plant cells due to the increased amounts of DNA, often leads to polyploid plants being larger in size than their diploid progenitors (Doyle and Coate, 2019; Bomblies, 2020; Clo and Kolář, 2021). This phenotypic change may increase their apparency to herbivores leading to a greater number of herbivores being found on polyploids. This change in size often comes at the cost of slower growth rate, which may in turn impact the plant’s ability to tolerate and recover from herbivore damage (Züst and Agrawal, 2016; Corneillie et al., 2018; DeRose et al., 2022; Anneberg et al., 2023a). Similarly, ploidy-driven changes in the photosynthetic rate of the plant could allow for faster or slower recovery depending on the direction (Warner and Edwards, 1993; Cao et al., 2018). Further, polyploidization may impact secondary-metabolite production, and polyploids may acquire higher levels of defenses against herbivores (Lavania et al., 2012; Bagheri and Mansouri, 2015; Edger et al., 2015; Gaynor et al., 2020).

While much effort has been focused on capturing the variation in the responses to polyploidy across different species, is also important to also recognize the intraspecific variation in response to polyploidy due to genetic differences in progenitor diploids (Soltis et al., 2016; Castro et al., 2020; Anneberg et al., 2023b; Bafort et al., 2023). Polyploidy can arise multiple times independently within a single taxon across genetically divergent individuals, leading to differences in whole-genome duplication’s affect within a single species (Soltis et al., 1993; Segraves et al., 1999). For example, in the context of species interactions, Anneberg et al. (2023a) recently found that the effect of polyploidy on duckweed microbiomes was different across multiple, independently synthesized neopolyploid lineages of duckweed. Yet, our knowledge of how other species interactions may be impacted by the interactions of polyploidization and genetic background, such as plant-herbivore interactions, is still limited.

Compared to other biotic interactions, evidence for polyploidy’s effect on plant-herbivore interactions are heavily weighted toward herbivore attraction, attack, and resultant plant performance, with none experimentally addressing the impact on both herbivore and plant populations (Arvanitis et al., 2010; Gross and Schiestl, 2015; Münzbergová et al., 2015; O’Connor et al., 2019). Additionally, results from these studies are mixed. For example, Halverson et. al (2008) found that neither diploid nor polyploid *Solidago altissima* were consistently attacked more frequently by five species of herbivores. Similarly, Thompson et al. (1997) found that the moth *Greya potentilla* was sometimes more likely to attack tetraploids over diploid *Heuchera grossularifolia*, but they were still able to colonize both diploids and polyploids of separate origin. Indeed, while insightful, general conclusions about the overall impact on the plant-herbivore relationship are hard to synthesize, as these studies are often conducted in a wide variety of habitat types, using a mixture of established and newly synthesized polyploids of different ages, and do not quantify herbivore or plant performance (Thompson et al., 1997; Walczyk and Hersch-Green, 2019; Harkin and Stewart, 2021; Harms and Walter, 2021). Notably, however, a recent study by Curé et al. (2022) did test the preference and performance of a specialist pea aphid, *Acyrthosiphon pisum*, on two different host plants. They found no ploidal-dependent preference in two species, red clover and alfalfa, but they did find that aphids that originated from populations specialized on diploid red clover had higher fecundity on that host than on synthesized neotetraploid red clover. To our knowledge, this is the only study to quantify herbivore performance on diploids and neopolyploids, and now opens the door to build on these results by also quantifying plant performance and expanding to new plant-herbivore systems.

While quantifying herbivory on natural established populations has provided key insights into the effects of polyploidy on plant-herbivore interactions, most prior studies suffer from three limitations. First, these can confound the impact of polyploidy, interspecific hybridization, and evolution following whole-genome duplication (Parisod et al., 2010; Drunen and Husband, 2018; Bomblies, 2020). By using newly established or synthetic polyploids, or neo-polyploids, one can isolate the immediate consequences of whole-genome duplication alone. Second, the outcome of whole-genome duplication may vary on the genotypic background of the individual, so multiple genotypes should be used, and genetic variance must be accounted for (Drunen and Husband, 2018; Wei et al., 2020; Bafort et al., 2023). Third, most studies of polyploidy-herbivory interactions are conducted over only a small portion of the life-history of the plant or herbivore and usually on single individuals. This may limit our ability to draw conclusions on the fitness impacts of polyploidy under herbivory. To our knowledge, there are no population-level experimental studies investigating both plant and herbivore performance and herbivore preference in the context of neopolyploidy. Curé et al. (2022) were the first and only to investigate this relationship using neopolyploids but focused on individual-level plant response. Consequently, to further our understanding of the outcomes of whole-genome duplication on species interactions, there is a need for experiments examining the immediate effects of whole-genome duplication at the population level, using multiple independently created polyploid genotypes (Parisod et al., 2010; Kolář et al., 2017; Spoelhof et al., 2017; Drunen and Husband, 2018; Anneberg et al., 2023b).

Duckweed is well suited to fill this knowledge gap in how neopolyploidy affects plant-herbivore interactions (Laird and Barks, 2018). Duckweeds are globally distributed, small, aquatic floating plants that primarily reproduce asexually via budding. An individual consists of a single frond, a small leaf-like structure making up the entire shoot, and multiple roots (Ziegler et al., 2015; Acosta et al., 2021). Duckweeds reproduce rapidly (within four days in optimal conditions), and thus multiple generations can be studied in the span of several weeks (Ziegler et al., 2015; Hart et al., 2019). While there is natural variation in many duckweed species, Greater duckweed (*S. polyrhiza*) is gaining traction as model system for polyploidy and herbivory owing to the amenability of population-level studies using synthesized neotetraploid plants, affording direct comparison of neopolyploid populations to those of their diploid progenitors (Anneberg et al., 2023a). Combined with experimental studies of herbivores, population-level impacts of herbivory can be precisely studied from both the plant and herbivore perspective (Mariani et al., 2020; Subramanian and Turcotte, 2020, 2023).

The water-lily aphid, *Rhopalosiphum nymphaeae*, is a globally-distributed generalist herbivore of duckweeds (Halder et al., 2020; Subramanian and Turcotte, 2020, 2023). Aphids are phloem-feeding herbivores that reproduce facultatively parthenogenetically via live birth with a population doubling time of around two days (Hance et al., 1994). Because both aphids and duckweeds are fast-reproducing, asexual organisms, together they provide a unique opportunity to evaluate the effect of neopolyploidy on population growth rates of both host and herbivore over multiple generations.

Here, we addressed four questions. 1) Do aphids exhibit a preference for diploid or neopolyploid duckweed, and if so, is this a function of the effect of ploidy on plant body size? 2) Does duckweed ploidy alter aphid population performance? 3 Do aphids differentially affect the performance of neopolyploid and diploid duckweed populations? 4) Are the results of the previous questions dependent upon the genetic lineage of duckweed?

## MATERIALS AND METHODS

### Cultivation of duckweed and aphids-

We used colchicine-induced autotetraploid and colchicine-exposed but unconverted diploids as described in Anneberg et al. (2023a) to answer our questions. To obtain these, we had previously applied the mitotic inhibitor colchicine to induce whole-genome duplication in six genetically distinct diploid *S. polyrhiza* collected from eastern Pennsylvania and western Ohio, U.S.A. (See Table S1 for collection site info) (Xu et al., 2018; Anneberg et al., 2023a; Kerstetter et al., 2023). Then, in 2019 and 2020, ploidy was confirmed using flow cytometry following Wei et al. (2020). Although we did not observe any residual effects of colchicine treatment, to be conservative, we used the colchicine-treated but unconverted diploid individuals in the experiment (Anneberg et al., 2023a). Prior to the experiment, we grew individual lineages of duckweed in the growth chamber at 23.5°C, 50% humidity, 50 μmol/m2 /s light, and 16:8 light/dark cycle.

Water-lily aphids were collected from a duckweed community composed of a mixture of several species of duckweed (*Spirodela polyrhiza, Lemna minor, and Wolffia brasiliensis*) at Twin Lakes Park in Westmoreland County, Pennsylvania, USA (40.323383333, -79.472383333) in September of 2017 (Subramanian and Turcotte, 2020). Stock aphid populations were then maintained on monocultures of diploid *S. polyrhiza* populations in the growth chamber.

### Preference Experiments-

To determine whether aphids exhibit feeding preference for diploid or neotetraploid duckweed, we conducted several two-way aphid choice trials following the basic set-up established in Subramanian and Turcotte (2020) in January and February of 2023. Each trial consisted of a diploid and its corresponding derived neopolyploid in preference arenas. On average, our neopolyploid duckweed were 46% larger in surface area than their diploid progenitors (See Supplemental Table 2). Considering this size difference, we conducted two separate sets of trials in preference arenas. The first was the ‘Abundance-controlled’ trial – we added exactly six fronds of each ploidy of a given lineage to the arenas. The second was the ‘Area-controlled’ trial – we added approximately equal population surface area of each ploidy of a given lineage to the preference arenas. We equalized ploidal surface area by first placing duckweed in 3.5 cm^2^ cells of a culture plate, such that there was a single, non-overlapping layer of duckweed floating on the surface before moving pairs of them to the preference arenas. The preference arenas consisted of 60 mL jars with 19.6 cm^2^ openings that were filled with 50 mL of 0.1x strength diluted, sterile plant growth media (Appenroth et al., 1996). In the center of the 60 mL jar, we floated a 0.6 cm diameter circle of white plastic as a platform for the aphid in the middle of the duckweed population. The diploid and neopolyploid plants were intermixed around the platform and then we placed a single 3^rd^ instar aphid on the platform. We tested preference on five of the six genetic lineage pairs of diploid-neopolyploid duckweeds. One genetic lineage (SP.07) was omitted due to contamination with algae that was later cleaned, allowing for use in the performance trials (see below). Both the Area- and Abundance-controlled trials were replicated 20 times per genetic lineage, for a total of 200 paired trials (5 genetic lineages x 2 trial types x 20 replicates). All aphids and duckweeds were only used once. We determined preference by observing which ploidy the aphid chose to insert its stylet (Subramanian and Turcotte, 2020). We recorded aphid preference after 1, 5, 30, 60 and 90 minutes and 24 hours. If the aphid died or crawled out of the jar (which only occurred 4 times out of 200 trials), no choice was recorded, and it was removed from the data analysis. For 93% of the trials, the aphid stayed on the same individual originally chosen (usually in the first 1 or 5 minutes), so only final choice at 24 hours was used in the analysis.

### Performance Experiment-

We assessed aphid and duckweed performance in a separate, full factorial experiment where we crossed ploidy, genetic lineage and aphid presence in a growth chamber. We added 220 mL of 0.5x strength growth media to 240 mL glass jars. Into each, we added six individuals of a single ploidy from a single genetic lineage of duckweed. We then randomly chose jars to add either five 3^rd^ instar aphids or no aphids. Each combination was replicated 10 times for a total of 240 jars (6 genetic lineages x 2 ploidies x 2 aphid treatments x 10 replicates). The 10 replicates were split into two time-blocks of five, run consecutively. Each experiment lasted 15 days, allowing for three-four generations of both duckweeds and aphids. We quantified population growth of the aphids by counting their abundances two-three times per week. We quantified duckweed performance in two ways, both of which represent multigenerational fitness. First, we quantified duckweed population abundance over time by counting their abundances two-three times per week. Because aphids and duckweed reproduce asexually under these conditions, abundance serves as a direct measure of population performance. Second, at the end of the 15 days, we measured final biomass by harvesting all duckweeds at the end of the experiment, drying them at 55°C for one week, and weighing them on Cahn C-31 microbalance scale to the nearest 0.0001 g.

### Statistical Analyses-

We tested for an effect of neopolyploidy on aphid preference, aphid performance and plant performance (in terms of both abundances and final biomass). All analyses were performed in R version 4.1.2 (R Core Team, 2021). For the preference trials, we conducted *G*-tests of goodness-of-fit for each lineage in each trial type. For each trial type (Area- or Abundance-controlled), we computed the total *G* (summed across lineages), pooled *G* and calculated the heterogeneity *G* to assess whether there was significant variation among genetic lineages using the *RVAideMemoire* package (Herve, 2023).

For the performance experiments, we constructed separate generalized linear models with aphid population growth, plant population growth and final plant biomass as response variables. For aphid population growth, we used a generalized linear mixed model (GLMM) with a negative binomial probability distribution, with aphid abundances as the response variable, and ploidy (diploid or neopolyploid) and genetic lineage (as a categorical factor) and their interaction as main effects. We also included experimental time-block as a fixed effect. Lastly, to account for repeated measures over the course of the experiment, ‘Day’, or day of sampling, and ‘Jar ID’, which was the individual experimental unit, were included as crossed random effects. We removed the abundance on day one from this analysis because all experimental units started with the same exact number of aphids (five). Linear mixed effects models were run using the *glmmTMB* package (Brooks et al., 2017). We used a negative binomial distribution to account for overdispersion in the aphid population data. All model residuals were assessed using the DHARMa package (Hartig, 2022).

For duckweed population abundances, we used a GLMM with a Poisson probability distribution with duckweed abundances as the response variable and similar explanatory variables and random effects as above, but we also included a main effect for herbivory (presence or absence) as well as a three-way interaction term (Ploidy:Lineage:Herbivory). We also removed the first day’s data point from this analysis because all experimental units started with the same exact number of duckweed fronds (six).

Finally, we analyzed final duckweed dry biomass using a linear model with a normal, Gaussian probability distribution with ploidy, genetic lineage and herbivory as main effects, with all interactions and time-block as a fixed effect.

We also ran lineage-specific GLMMs for all models (aphid abundance, duckweed abundance and duckweed biomass) post-hoc to see which, if any, genetic lineages were driving significant effects. Lineage-specific models had the same structure as the overall models but without the ‘genetic lineage’ response variable.

## RESULTS

### Herbivore preference and performance-

When duckweed ploidies were matched by frond abundance in the Abundance-controlled trial, we found a significant aphid preference for neopolyploid plants over diploids (Fig. 1A, Table 1). Pooled across all lineages, aphids chose polyploids 66 out of 100 trials, and this preference was consistent across all genetic lineage pairs. Given that diploids were only chosen 31 times implies that neopolyploids were 213% as likely to be attacked by aphids than diploids. However, when duckweed ploidies were matched by total surface area in the Area-controlled trial, aphids did not exhibit significant preference (Fig. 1B, Table 1). Specifically, in 54 of the trials, the aphid chose the neopolyploid, and in 46 of the trials, the aphid chose the diploid. This was consistent across lineages (Table 2).

**Figure 1.**
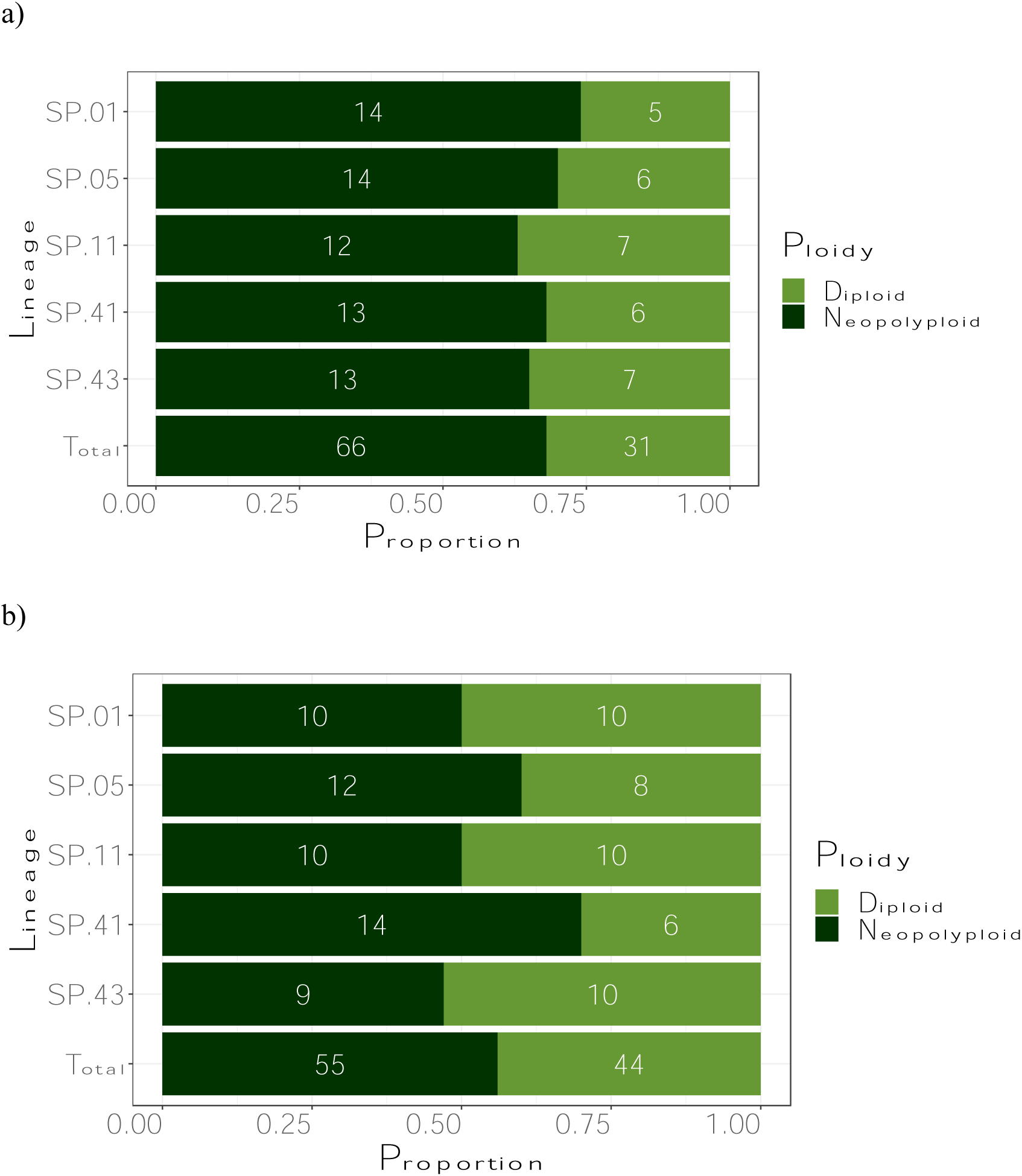
Water-lily aphid herbivore preference (number of trials aphids chose diploid or neopolyploid plant) in the **a)** Abundance-controlled trial and the **b)** Area-controlled trial. Results are presented by lineage (number) and summed across all lineages (Total).

**Table 1.** Preference trial results and G-values in the Abundance controlled trial (when *equal number of duckweed (Spirodela polyrhiza) fronds* were used). G-values, degrees of freedom, and p-values are given for each lineage, their sum, as well as pooled across all lineages.

| Lineage | Diploid<br>chosen | Neopolyploid<br>chosen |  | G-value | d.f. | P-value |
| --- | --- | --- | --- | --- | --- | --- |
| SP.01 | 5 | 14 |  | 4.43 | 1 | 0.035 |
| SP.05 | 6 | 14 |  | 3.29 | 1 | 0.069 |
| SP.11 | 7 | 12 |  | 1.33 | 1 | 0.25 |
| SP.41 | 6 | 13 |  | 2.64 | 1 | 0.10 |
| SP.43 | 7 | 13 |  | 1.83 | 1 | 0.18 |
|  |  |  | total G | 13.53 | 5 | 0.019 |
| pooled | 31 | 66 | pooled G | 12.92 | 1 | 0.00033 |
|  |  |  | heterogeneity G | 0.6121 | 4 | 0.96 |

**Table 2.** Preference trial results and G-values in the Area Controlled trial (when *equal total area* of duckweeds were used). G-values, degrees of freedom, and p-values are given for each lineage, their sum, as well as pooled across all lineages.

| Lineage | Diploid<br>chosen | Neopolyploid<br>chosen |  | G-value | d.f. | P-value |
| --- | --- | --- | --- | --- | --- | --- |
| SP.01 | 10 | 10 |  | 0 | 1 | 1 |
| SP.05 | 8 | 12 |  | 0.81 | 1 | 0.82 |
| SP.11 | 10 | 10 |  | 0 | 1 | 1 |
| SP.41 | 6 | 14 |  | 3.29 | 1 | 0.07 |
| SP.43 | 10 | 9 |  | 0.05 | 1 | 0.82 |
|  |  |  | total G | 4.15 | 5 | 0.53 |
| pooled | 44 | 55 | pooled G | 1.22 | 1 | 0.27 |
|  |  |  | heterogeneity G | 2.92 | 4 | 0.57 |

Plant polyploidy alone increased aphid performance but genetic lineage and its interaction with ploidy also significantly affected aphid performance (P < 0.001, Fig. 2, Table S3). Neopolyploids, on average, hosted 14% more aphids than diploids at the end of the experiment. The significant ploidy-lineage interaction indicates that the effect of ploidy on performance varied by lineage; for example, neopolyploid SP.11 hosted an average of 33% *more* aphids than diploid SP.11 by the end of the experiment, whereas aphids on SP.07 performed 4% worse on polyploids than the diploid (Ploidy:Lineage interaction, df = 5, P < 0.001, Fig. 2, Table S3). Only one lineage, SP.11, showed significant effects of neopolyploidy on aphid abundance in the lineage-specific models (Table S4).

**Figure 2.**
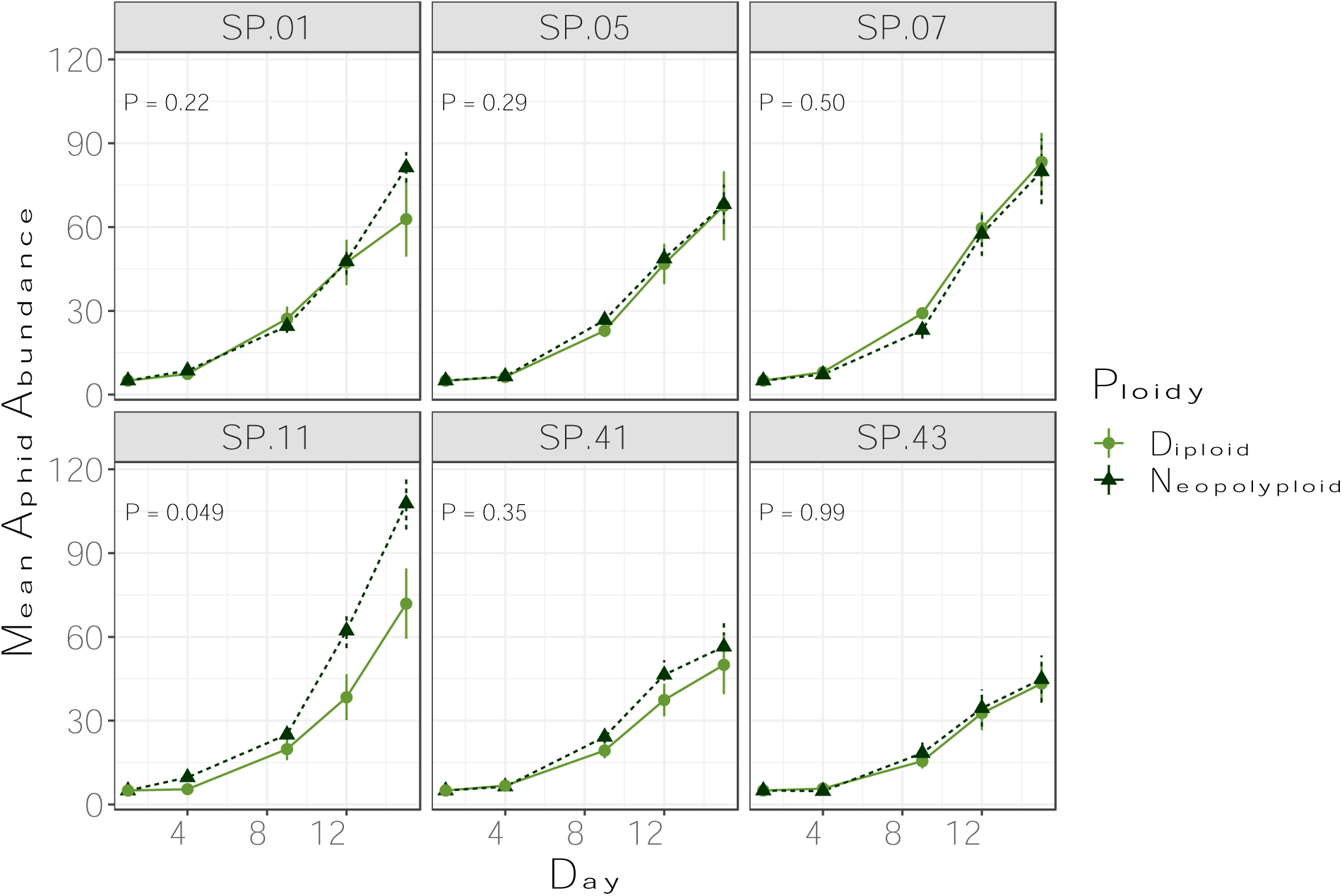
Aphid abundances over time when feeding on diploid or neopolyploid duckweed. Each panel represents a different duckweed lineage, and each point is the mean and standard error of 10 replicates. P-values on the graph represent the significance of ploidy in the lineage-specific GLMMs (See Table S4 for full model results)

### Duckweed Performance-

#### Abundance-

Overall, neopolyploid duckweed reached lower abundances in the face of herbivory than diploid duckweed in a lineage-dependent manner (three-way Ploidy:Herbivory:Lineage interaction, P = 0.031, Fig. 3, Table S5). For most lineages, neopolyploid duckweed abundance was more impacted by herbivory than diploid abundance, but the result was highly dependent on the genetic lineage. Independent duckweed lineages also responded to neopolyploidy differently (Ploidy:Lineage interaction, df = 5, P < 0.001, Fig. 3, Table S5). And while there was no Ploidy:Herbivory interaction (neopolyploids *overall* were not significantly more or less tolerant than diploids in terms of abundance), there were strong lineage-dependent responses to herbivory

**Figure 3.**
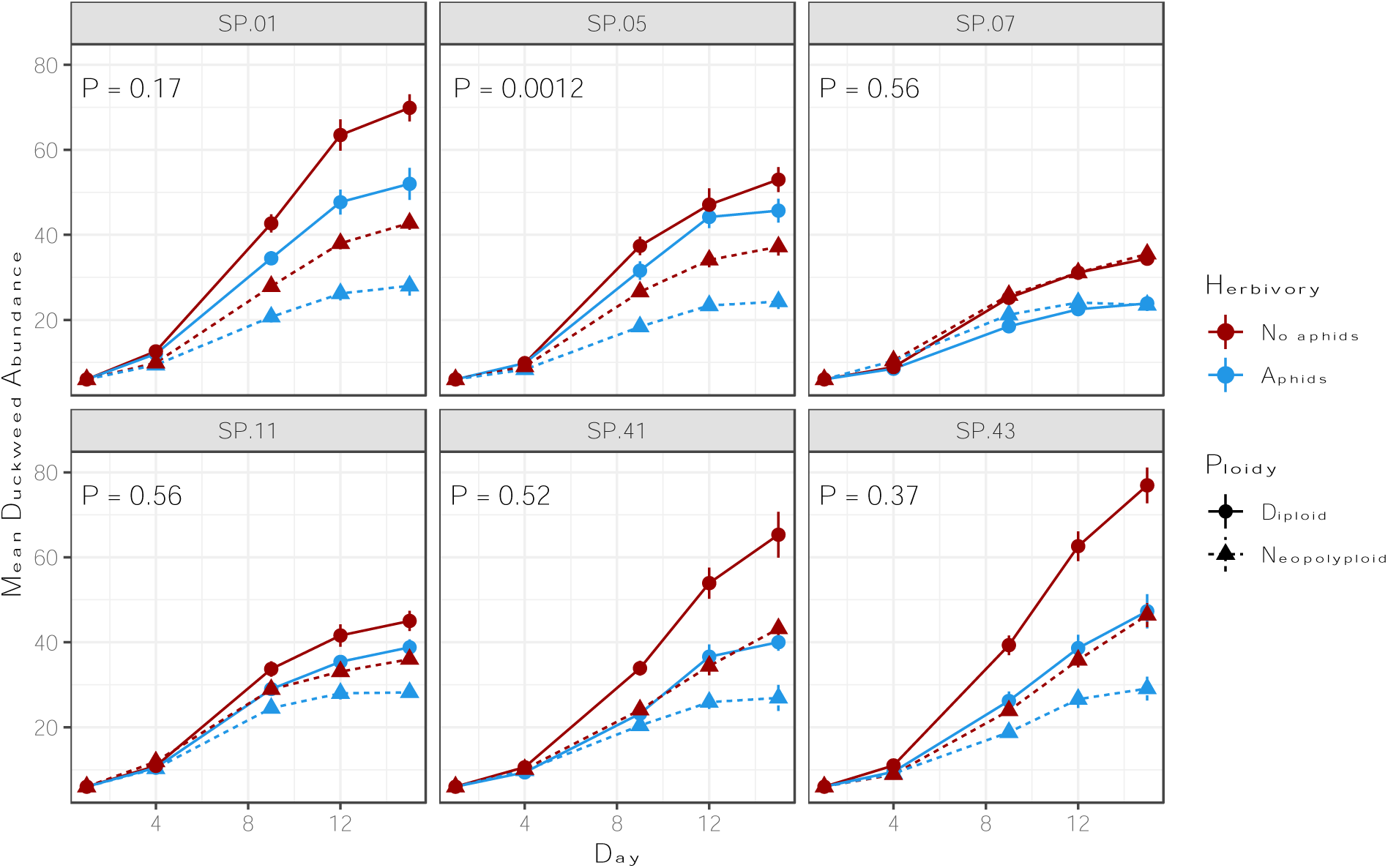
Duckweed abundance over time for diploid and neopolyploids with (red) and without aphids (blue). Each panel represents a different duckweed lineage, and each point represents the mean and standard error of 10 replicates. P values represent the Ploidy:Herbivory interaction and are calculated from the lineage-specific Poisson GLMMs (See Table S6 for full model results).

(Lineage:Herbivory interaction, df = 5, P <0.001, Fig 3, Table S5). For example, SP.01 was very tolerant, with its average final abundance only decreased by 11% (across diploids and neopolyploids) in the face of herbivory, whereas SP.43 declined by 38% suggesting lower tolerance. It is worth noting, however, that lineage-specific models reveal that SP.05 was the only lineage to exhibit a significant Ploidy:Herbivory interaction, wherein the neopolyploid’s abundance was more significantly impacted and less tolerant to aphids than the diploid (Table S6). Diploid SP.05 final abundance declined by an average of 14% in the face of herbivory, whereas neopolyploid’s final abundance declined by an average 35%.

#### Biomass-

Aphid presence significantly reduced duckweed final biomass across all treatments but did so unevenly among ploidies (Fig. 4, Table S7, Table S8). Aphids impacted neopolyploid biomass similarly to diploid biomass (2% difference in their tolerance) at a marginally significant level (Fig. 4, Table S7, Ploidy:Herbivory interaction, df = 1, *P* = 0.063), but the biological effect size was very small (Table S7). This result, however, varied among genetic lineages at a marginally significant level (Ploidy:Herbivory:Lineage interaction, df = 5, *P* = 0.065); neopolyploids of SP.05, SP.11 and SP.41 were all less tolerant than their diploid progenitors in terms of biomass, whereas diploids of SP.01, SP.07 and SP.43 were all less tolerant than neopolyploids.

**Figure 4.**
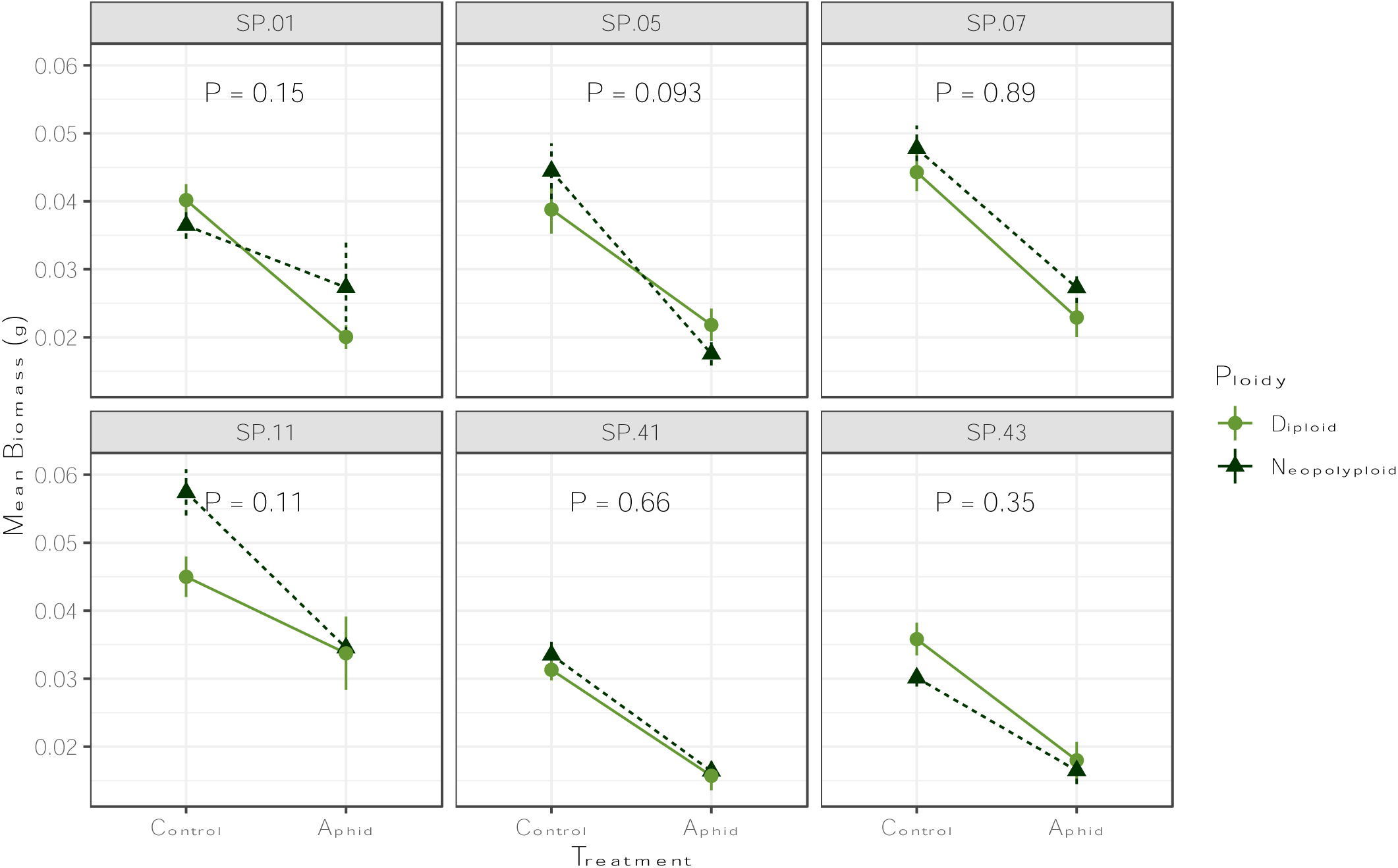
Duckweed biomass by genotype: Genotype-specific changes in biomass following the addition of herbivory in both the diploid and the polyploid treatments.

## DISCUSSION

Our results demonstrated significant differences in the effect of polyploidy and genetic lineage on herbivore preference and performance, and plant performance in response to herbivory. Preference experiments indicated that aphids preferred neopolyploid duckweed across all genetic lineages, but that this result was largely driven by differences between ploidies in frond size, as the preference disappeared when we controlled for size. In addition to aphids preferring the neopolyploid duckweed, aphids also often performed better -- reaching higher abundances--on neopolyploid duckweed. However, this result was highly dependent on plant genetic lineage. Lastly, neopolyploids appeared to be slightly less tolerant than diploids in the face of herbivory, but the effects were small and highly dependent on genetic lineage.

The relationships between herbivory, polyploidy, and plant genetic background on plant performance were complex. The absence of a significant two-way interaction between polyploidy and herbivory on plant performance, but the presence of three-way interactions, suggests that the effects of polyploidy and herbivory do not depend on each other in a straightforward manner, and that the genetic background of the plant plays a large role. For example, in the lineage SP.05, aphids reached similar abundances on neopolyploids and diploids, but abundance and biomass of neopolyploid SP.05 were more negatively impacted by aphid herbivory than its diploid progenitor (Fig. 3, Fig. 4, Table S6, Table S8.). This would imply that neopolyploid duckweed are, in fact, less tolerant per herbivore. However, neopolyploid SP.11 hosted more aphids than diploid SP.11, but there were no significant differences in how they responded to herbivory. This may actually imply that neopolyploid SP.11 is *more* tolerant per aphid than diploid SP.11. Such findings emphasize the need for a nuanced understanding of how multiple factors interact to shape ecological outcomes. Had we only quantified herbivore performance or plant performance, these complex relationships could have been overlooked. However, to truly make comparisons and broad conclusions across multiple plant-herbivore systems, more population-level data on the plant and the herbivore in the context of polyploidy are needed (Curé et al., 2022; Anneberg et al., 2023b).

Overall, aphids preferred neopolyploids and performed better on them. Indeed, this trend has been found in other plant herbivore systems with mixed ploidies, such as *Greya politella* moths and plants in the *Lithophragma* genus, and Alligator weed (*Alternanthera philoxeroides*) and the flea beetle (*Agasicles hygrophila*) (Janz and Thompson, 2002; Krug and Sosa, 2019). However, many studies reporting this pattern also confirm that their results depended on other factors, such as plant origin, year sampled, and environmental context, and the opposite trend has also been found in some plant-herbivore systems as well (Janz and Thompson, 2002; Hull-Sanders et al., 2009; König et al., 2014, 2016; Münzbergová et al., 2015). Our results with synthetic neopolyploids show that, in particular, it is important to account for the differences in size between the two ploidies as a potential mechanism driving herbivore preference, yet this control is not commonly conducted. Corroborating the preference study result that plant size plays a large role in driving these interactions, in the performance study, the lineage with the smallest size difference between diploids and neopolyploids, SP.07, exhibited very little differences in aphid population sizes (See Supplemental Table 2). Similarly, SP.11 neopolyploids saw the largest increase in aphid abundances compared to diploids, and they are approximately 56% larger than their diploid progenitors. However, this trend was not always consistent across lineages; neopolyploid SP.05 are nearly double the size of their diploid progenitors, but they exhibited little differences in the number of aphids hosted. Interestingly, this may indicate that something other than size may be contributing to polyploidizations effect on the duckweed-aphid relationship for this lineage. There is very little mechanistic work behind what might drive differences between herbivore preference and performance *in the context of polyploidy*. However, we do know that, in addition to ‘apparency’ can be driven by visual cues such as size, preferences are also driven by olfactory, mechanical, and chemical cues, and that variation in plant chemistry exists even at the intraspecific level (Powell et al., 2006; Jakobs and Müller, 2018; Endara et al., 2023). For example, in *Centaurea phyrgia*, Münzbergovà et al. (2015) cite differences in levels of secondary compounds gallic acid and several polyphenols in diploid *Centaurea phyrgia*, as a potential mechanism for diploids suffering less seed damage than polyploids. Further, polyploidization may have varied effects on herbivores with alternative modes of feeding, such as chewing or mining, as compared to phloem-suckers like aphids.

Feeding modes vary amount and type of damage they inflict on host plants, and thus plants may use different strategies to tolerate and resist different herbivores (Ali and Agrawal, 2012; Marmolejo et al., 2021; Xu et al., 2021). How chemical versus structural defenses change with whole-genome duplication, however, remains unknown. Thus, one avenue for future research could involve investigating the chemical profiles of neopolyploid and diploid duckweed and whether these impact varying types of herbivory differently.

## CONCLUSIONS

Our experiment, using multiple neopolyploid lineages, revealed that that polyploidy and genetic lineage impact herbivore preferences for plant hosts and herbivore fitness, and that this trend is, in part, driven by size differences between neopolyploids and diploids. This study represents one of the initial attempts to investigate these dynamics in aquatic plants, and further work should consider expanding its scope to encompass a broader range of aquatic species. The extent to which neopolyploids’ fitness is affected by herbivory was strongly dependent on genetic lineage. By combining results on herbivore performance and plant performance across multiple plant genotypes, we were able to uncover complex relationships between ploidy, herbivory, plant genetic background. Our work here lays the groundwork for future experimental studies to explore the longer-term and mechanistic drivers of the impact of polyploidy on plant-herbivore relationships, as well as to further understand how these impacts might impact plant and herbivore establishment and persistence in nature.

### Future directions-

Long-term, polyploidy can alter species interactions in two ways: either via the direct changes caused by whole-genome duplication, or by the indirect changes incurred via evolution that occurs following whole-genome duplication. Here, we were able to investigate the effects of the direct changes caused by polyploidization, leaving the latter still up for investigation. A recent study by Malacrinò et al. (2022), however, showed that exposure to herbivory rapidly increased *S. polyrhiza* resistance in only 30 generations. Future research using long-term experimental studies with both neopolyploids and diploids would be needed to address whether polyploids would evolve differently than diploids in the presence/absence of herbivores.

Given our results, it is possible that, in addition to facilitating establishment of polyploids in natural communities, whole-genome duplication in plants may also contribute to the evolutionary diversification of the herbivore. Aphids not only incorporated neopolyploid duckweed in their diets, but also preferred neopolyploids and performed better on them; this implies that neopolyploidy may be a mechanism to advance the migration of herbivores and facilitate aphid expansion outside of their current ranges (Curé et al., 2022).

## Acknowledgements

The authors thank Elizabeth O’Neill for synthesizing the neopolyploids, confirming ploidy using flow cytometry and maintaining the *S. polyrhiza* stock populations. We thank Jae Kerstetter and Audrey Burr for assistance in maintaining the *S. polyrhiza* genotypes and Jae Kerstetter, Lacey Rzodkiewicz and Cara Faillace for assistance in maintaining the *R. nymphaeae* stocks. The authors also thank Dr. Thomas Anneberg and Jason Simmons for their help carrying out the project. Lastly, the authors thank the Dietrich School of Arts and Sciences and grants from the National Science Foundation (NSF-GRFP to HRA, DEB-2027604 to T-LA, DEB-1935410 to MMT) for funding this research.

## Author Contributions

HRA, T-LA and MMT conceptualized and designed the study. HRA carried out the experiments. HRA, T-LA and MMT conceptualized the analysis. HRA analyzed the data and wrote the first draft of the manuscript, and HRA, T-LA and MMT edited subsequent drafts.

## Data Availability Statement

Data and R scripts will be archived in an online repository by the time of publication.

## Supporting Information

Additional supporting information may be found online in the Supporting Information section at the end of the article.

## Literature Cited

Acosta, K., K. J. Appenroth, L. Borisjuk, M. Edelman, U. Heinig, M. A. K. Jansen, T. Oyama, et al. 2021. Return of the Lemnaceae: duckweed as a model plant system in the genomics and postgenomics era. The Plant Cell 33: 3207–3234.

Ali, J. G., and A. A. Agrawal. 2012. Specialist versus generalist insect herbivores and plant defense. Trends in Plant Science 17: 293–302.

Anneberg, T. J., E. M. O’Neill, T.-L. Ashman, and M. M. Turcotte. 2023a. Polyploidy impacts population growth and competition with diploids: multigenerational experiments reveal key life history tradeoffs. New Phytologist.

Anneberg, T. J., M. M. Turcotte, and T.-L. Ashman. 2023b. Plant neopolyploidy and genetic background differentiates the microbiome of duckweed across a variety of natural freshwater sources. bioRxiv: 2023.04.29.538806.

Appenroth, K.-J., S. Teller, and M. Horn. 1996. Photophysiology of turion formation and germination inSpirodela polyrhiza. Biologia Plantarum 38: 95–106.

Arvanitis, L., C. Wiklund, Z. Münzbergova, J. P. Dahlgren, and J. Ehrlén. 2010. Novel antagonistic interactions associated with plant polyploidization influence trait selection and habitat preference. Ecology Letters 13: 330–337.

Bafort, Q., T. Wu, A. Natran, O. D. Clerck, and Y. V. de Peer. 2023. The immediate effects of polyploidization of Spirodela polyrhiza change in a strain-specific way along environmental gradients. Evolution Letters.

Bagheri, M., and H. Mansouri. 2015. Effect of Induced Polyploidy on Some Biochemical Parameters in Cannabis sativa L. Applied Biochemistry and Biotechnology 175: 2366–2375.

Bomblies, K. 2020. When everything changes at once: finding a new normal after genome duplication. Proceedings of the Royal Society B: Biological Sciences 287: 20202154.

Brooks, M., K. Kristensen, K. van Benthem, A. Magnusson, C. Berg, A. Nielsen, H. Skaug, et al. 2017. glmmTMB Balances Speed and Flexibility Among Packages for Zero-inflated Generalized Linear Mixed Modeling. The R Journal 2: 378–400.

Cao, Q., X. Zhang, X. Gao, L. Wang, and G. Jia. 2018. Effects of ploidy level on the cellular, photochemical and photosynthetic characteristics in Lilium FO hybrids. Plant Physiology and Biochemistry 133: 50–56.

Castro, M., J. Loureiro, A. Figueiredo, M. Serrano, B. C. Husband, and S. Castro. 2020. Different Patterns of Ecological Divergence Between Two Tetraploids and Their Diploid Counterpart in a Parapatric Linear Coastal Distribution Polyploid Complex. Frontiers in Plant Science 11: 315.

Clo, J., and F. Kolář. 2021. Short- and long-term consequences of genome doubling: a meta- analysis. American Journal of Botany 108: 2315–2322.

Comai, L. 2005. The advantages and disadvantages of being polyploid. Nature Reviews Genetics 6: 836–846.

Corneillie, S., N. D. Storme, R. V. Acker, J. U. Fangel, M. D. Bruyne, R. D. Rycke, D. Geelen, et al. 2018. Polyploidy Affects Plant Growth and Alters Cell Wall Composition . Plant Physiology 179: 74–87.

Curé, A. E., D. M. Althoff, and K. A. Segraves. 2022. Host expansion in a specialist herbivore is facilitated by whole-genome duplication in the host plant. Ecological Entomology.

DeRose, R. J., R. S. Gardner, R. L. Lindroth, and K. E. Mock. 2022. Polyploidy and growth— defense tradeoffs in natural populations of western quaking Aspen. Journal of Chemical Ecology: 1–10.

Doyle, J. J., and J. E. Coate. 2019. Polyploidy, the Nucleotype, and Novelty: The Impact of Genome Doubling on the Biology of the Cell. International Journal of Plant Sciences 180: 1– 52.

Drunen, W. E. V., and B. C. Husband. 2018. Immediate vs. evolutionary consequences of polyploidy on clonal reproduction in an autopolyploid plant. Annals of Botany 122: 195–205.

Edger, P. P., H. M. Heidel-Fischer, M. Bekaert, J. Rota, G. Glöckner, A. E. Platts, D. G. Heckel, et al. 2015. The butterfly plant arms-race escalated by gene and genome duplications. Proceedings of the National Academy of Sciences 112: 8362–8366.

Endara, M.-J., D. L. Forrister, and P. D. Coley. 2023. The Evolutionary Ecology of Plant Chemical Defenses: From Molecules to Communities. *Annual Review of Ecology*, Evolution, and Systematics 54.

Forrester, N. J., and T.-L. Ashman. 2017. The direct effects of plant polyploidy on the legume– rhizobia mutualism. Annals of Botany 121: 209–220.

Forrester, N. J., M. Rebolleda-Gómez, J. L. Sachs, and T.-L. Ashman. 2020. Polyploid plants obtain greater fitness benefits from a nutrient acquisition mutualism. New Phytologist 227: 944–954.

Fox, D. T., D. E. Soltis, P. S. Soltis, T.-L. Ashman, and Y. V. de Peer. 2020. Polyploidy: A Biological Force From Cells to Ecosystems. Trends in Cell Biology 30: 688–694.

Gaynor, M. L., S. Lim-Hing, and C. M. Mason. 2020. Impact of genome duplication on secondary metabolite composition in non-cultivated species: A systematic meta-analysis. Annals of Botany 126: mcaa107-.

Godfree, R. C., D. J. Marshall, A. G. Young, C. H. Miller, and S. Mathews. 2017. Empirical evidence of fixed and homeostatic patterns of polyploid advantage in a keystone grass exposed to drought and heat stress. Royal Society Open Science 4: 170934.

Gross, K., and F. P. Schiestl. 2015. Are tetraploids more successful? Floral signals, reproductive success and floral isolation in mixed-ploidy populations of a terrestrial orchid. Annals of Botany 115: 263–273.

Halder, J., A. B. Rai, S. Chakrabarti, and D. Dey. 2020. Distribution, Host Range and Bionomics of Rhopalosiphum nymphaeae (Linnaeus, 1761) a Polyphagous Aphid in Aquatic Vegetables. Defence Life Science Journal 5: 49–53.

Hamarashid, S. H., Y. Khaledian, and F. Soleimani. 2022. In vitro polyploidy-mediated enhancement of secondary metabolites content in Stachys byzantina L. Genetic Resources and Crop Evolution 69: 719–728.

Hance, Th., D. Nibelle, Ph. Lebrun, G. Impe, and C. Hove. 1994. Selection of Azolla forms resistant to the water lily aphid, Rhopalosiphum nymphaeaeLife history of Rhopalosiphum nymphaeae. Entomologia Experimentalis et Applicata 70: 11–17.

Harkin, C., and A. J. A. Stewart. 2021. Differential outcomes of novel plant-herbivore associations between an invading planthopper and native and invasive Spartina cordgrass species. Oecologia 195: 983–994.

Harms, N. E., and D. J. Walter. 2021. Influence of Butomus umbellatus L. lineage and age on leaf chemistry and performance of a generalist caterpillar. Aquatic Botany 172: 103391.

Hart, S. P., M. M. Turcotte, and J. M. Levine. 2019. Effects of rapid evolution on species coexistence. Proceedings of the National Academy of Sciences 116: 201816298.

Hartig, F. 2022. DHARMa: Residual Diagnostics for Hierarchical (Multi-Level / Mixed) Regression Models. R package version 0.4.6.

Herve, M. 2023. RVAideMemoire: Testing and Plotting Procedures for Biostatistics. R package version 0.9–83.

Hull-Sanders, H. M., R. H. Johnson, H. A. Owen, and G. A. Meyer. 2009. Influence of polyploidy on insect herbivores of native and invasive genotypes of Solidago gigantea (Asteraceae). Plant Signaling & Behavior 4: 893–895.

Jakobs, R., and C. Müller. 2018. Effects of intraspecific and intra-individual differences in plant quality on preference and performance of monophagous aphid species. Oecologia 186: 173– 184.

Janz, N., and J. N. Thompson. 2002. Plant polyploidy and host expansion in an insect herbivore. Oecologia 130: 570–575.

Jiao, Y., N. J. Wickett, S. Ayyampalayam, A. S. Chanderbali, L. Landherr, P. E. Ralph, L. P. Tomsho, et al. 2011. Ancestral polyploidy in seed plants and angiosperms. Nature 473: 97– 100.

Kerstetter, J. E., A. L. Reid, J. T. Armstrong, T. A. Zallek, T. T. Hobble, and M. M. Turcotte. 2023. Characterization of microsatellite markers for the duckweed Spirodela polyrhiza and Lemna minor tested on samples from Europe or the United States of America. bioRxiv: 2023.02.15.528655.

Kolář, F., M. Čertner, J. Suda, P. Schönswetter, and B. C. Husband. 2017. Mixed-Ploidy Species: Progress and Opportunities in Polyploid Research. Trends in Plant Science 22: 1041–1055.

König, M. A. E., C. Wiklund, and J. Ehrlén. 2016. Butterfly oviposition preference is not related to larval performance on a polyploid herb. Ecology and Evolution 6: 2781–2789.

König, M. A. E., C. Wiklund, and J. Ehrlén. 2014. Context-dependent resistance against butterfly herbivory in a polyploid herb. Oecologia 174: 1265–1272.

Krug, P., and A. J. Sosa. 2019. Mother knows best: plant polyploidy affects feeding and oviposition preference of the alligator weed biological control agent, Agasicles hygrophila. BioControl 64: 623–632.

Laird, R. A., and P. M. Barks. 2018. Skimming the surface: duckweed as a model system in ecology and evolution. American Journal of Botany 105: 1962–1966.

Lavania, U. C., S. Srivastava, S. Lavania, S. Basu, N. K. Misra, and Y. Mukai. 2012. Autopolyploidy differentially influences body size in plants, but facilitates enhanced accumulation of secondary metabolites, causing increased cytosine methylation. The Plant Journal 71: 539–549.

Levin, D. A., and D. E. Soltis. 2018. Factors promoting polyploid persistence and diversification and limiting diploid speciation during the K–Pg interlude. Current Opinion in Plant Biology 42: 1–7.

Madlung, A. 2013. Polyploidy and its effect on evolutionary success: old questions revisited with new tools. Heredity 110: 99–104.

Magalhães, T. L., K. Murphy, A. Efremov, V. Chepinoga, T. A. Davidson, and E. Molina-Navarro. 2021. Ploidy state of aquatic macrophytes: Global distribution and drivers. Aquatic Botany 173: 103417.

Malacrinò, A., L. Böttner, S. Nouere, M. Huber, M. Schäfer, and S. Xu. 2022. Induced responses contribute to rapid plant adaptation to herbivory. bioRxiv: 2022.11.24.517793.

Mariani, F., A. D. Giulio, S. Fattorini, and S. Ceschin. 2020. Experimental evidence of the consumption of the invasive alien duckweed Lemna minuta by herbivorous larvae of the moth Cataclysta lemnata in Italy. Aquatic Botany 161: 103172.

Marmolejo, L. O., M. N. Thompson, and A. M. Helms. 2021. Defense Suppression through Interplant Communication Depends on the Attacking Herbivore Species. Journal of Chemical Ecology 47: 1049–1061.

McCarthy, E. W., M. W. Chase, S. Knapp, A. Litt, A. R. Leitch, and S. C. L. Comber. 2016. Transgressive phenotypes and generalist pollination in the floral evolution of Nicotiana polyploids. Nature Plants 2: 16119.

Münzbergová, Z., J. Skuhrovec, and P. Maršík. 2015. Large differences in the composition of herbivore communities and seed damage in diploid and autotetraploid plant species. Biological Journal of the Linnean Society 115: 270–287.

Nuismer, S. L., and J. N. Thompson. 2001. Plant polyploidy and non-uniform effects on insect herbivores. Proceedings of the Royal Society of London. Series B: Biological Sciences 268: 1937–1940.

O’Connor, T. K., R. G. Laport, and N. K. Whiteman. 2019. Polyploidy in creosote bush (Larrea tridentata) shapes the biogeography of specialist herbivores. Journal of Biogeography 46: 597–610.

Parisod, C., R. Holderegger, and C. Brochmann. 2010. Evolutionary consequences of autopolyploidy. New Phytologist 186: 5–17.

Porturas, L. D., T. J. Anneberg, A. E. Curé, S. Wang, D. M. Althoff, and K. A. Segraves. 2019. A meta-analysis of whole genome duplication and the effects on flowering traits in plants. American Journal of Botany 106: 469–476.

Powell, G., C. R. Tosh, and J. Hardie. 2006. Host plant selection by aphids: Behavioral, evolutionary, and applied perspectives. Annual Review of Entomology 51: 309–330.

Ramsey, J., and T. S. Ramsey. 2014. Ecological studies of polyploidy in the 100 years following its discovery. Philosophical Transactions of the Royal Society B: Biological Sciences 369: 20130352.

R Core Team. 2021. R: A language and environment for statistical computing. R Foundation for Statistical Computing, Vienna, Austria.

Rezende, L., J. Suzigan, F. W. Amorim, and A. P. Moraes. 2020. Can plant hybridization and polyploidy lead to pollinator shift? Acta Botanica Brasilica 34: 229–242.

Rice, A., P. Šmarda, M. Novosolov, M. Drori, L. Glick, N. Sabath, S. Meiri, et al. 2019. The global biogeography of polyploid plants. Nature Ecology & Evolution 3: 265–273.

Segraves, K. A., and T. J. Anneberg. 2016. Species interactions and plant polyploidy. American Journal of Botany 103: 1326–1335.

Segraves, K. A., J. N. Thompson, P. S. Soltis, and D. E. Soltis. 1999. Multiple origins of polyploidy and the geographic structure of Heuchera grossulariifolia. Molecular Ecology 8: 253–262.

Soltis, D. E., V. A. Albert, J. Leebens-Mack, C. D. Bell, A. H. Paterson, C. Zheng, D. Sankoff, et al. 2009. Polyploidy and angiosperm diversification. American Journal of Botany 96: 336– 348.

Soltis, D. E., P. S. Soltis, and L. H. Rieseberg. 1993. Molecular Data and the Dynamic Nature of Polyploidy. Critical Reviews in Plant Sciences 12: 243–273.

Soltis, D. E., C. J. Visger, D. B. Marchant, and P. S. Soltis. 2016. Polyploidy: Pitfalls and paths to a paradigm. American Journal of Botany 103: 1146–1166.

Song, X.-M., J.-P. Wang, P.-C. Sun, X. Ma, Q.-H. Yang, J.-J. Hu, S.-R. Sun, et al. 2020. Preferential gene retention increases the robustness of cold regulation in Brassicaceae and other plants after polyploidization. Horticulture Research 7: 20.

Spoelhof, J. P., P. S. Soltis, and D. E. Soltis. 2017. Pure polyploidy: Closing the gaps in autopolyploid research. Journal of Systematics and Evolution 55: 340–352.

Subramanian, S. K., and M. M. Turcotte. 2023. Experimentally quantifying impact of herbivory on duckweed communities in natural pond ecosystems.

Subramanian, S. K., and M. M. Turcotte. 2020. Preference, performance, and impact of the water-lily aphid on multiple species of duckweed. Ecological Entomology 45: 1466–1475.

Thompson, J. N., B. M. Cunningham, K. A. Segraves, D. M. Althoff, and D. Wagner. 1997. Plant Polyploidy and Insect/Plant Interactions. The American Naturalist 150: 730–743.

Thompson, J. N., S. L. Nusimer, and K. Merg. 2004. Plant polyploidy and the evolutionary ecology of plant/animal interactions. Biological Journal of the Linnean Society 82: 511–519.

Tossi, V. E., L. J. M. Tosar, L. E. Laino, J. Iannicelli, J. J. Regalado, A. S. Escandón, I. Baroli, et al. 2022. Impact of polyploidy on plant tolerance to abiotic and biotic stresses. Frontiers in Plant Science 13: 869423.

Van-de-Peer, Y., T.-L. Ashman, P. S. Soltis, and D. E. Soltis. 2020. Polyploidy: an evolutionary and ecological force in stressful times. The Plant Cell 33: 11–26.

Walczyk, A. M., and E. I. Hersch-Green. 2019. Impacts of soil nitrogen and phosphorus levels on cytotype performance of the circumboreal herb Chamerion angustifolium: implications for polyploid establishment. American Journal of Botany 106: 906–921.

Warner, D. A., and G. E. Edwards. 1993. Effects of polyploidy on photosynthesis. Photosynthesis Research 35: 135–147.

Wei, N., R. Cronn, A. Liston, and T.-L. Ashman. 2019. Functional trait divergence and trait plasticity confer polyploid advantage in heterogeneous environments. The New Phytologist 221: 2286–2297.

Wei, N., Z. Du, A. Liston, and T.-L. Ashman. 2020. Genome duplication effects on functional traits and fitness are genetic context and species dependent: studies of synthetic polyploid Fragaria. American Journal of Botany 107: 262–272.

Wood, T. E., N. Takebayashi, M. S. Barker, I. Mayrose, P. B. Greenspoon, and L. H. Rieseberg. 2009. The frequency of polyploid speciation in vascular plants. Proceedings of the National Academy of Sciences 106: 13875–13879.

Xu, J., X. Wang, H. Zu, X. Zeng, I. T. Baldwin, Y. Lou, and R. Li. 2021. Molecular dissection of rice phytohormone signaling involved in resistance to a piercing-sucking herbivore. New Phytologist 230: 1639–1652.

Xu, N., F. Hu, J. Wu, W. Zhang, M. Wang, M. Zhu, and J. Ke. 2018. Characterization of 19 polymorphic SSR markers in Spirodela polyrhiza (Lemnaceae) and cross-amplification in Lemna perpusilla. Applications in Plant Sciences 6: e01153.

Yang, P.-M., Q.-C. Huang, G.-Y. Qin, S.-P. Zhao, and J.-G. Zhou. 2014. Different drought-stress responses in photosynthesis and reactive oxygen metabolism between autotetraploid and diploid rice. Photosynthetica 52: 193–202.

Ziegler, P., K. Adelmann, S. Zimmer, C. Schmidt, and K. -J. Appenroth. 2015. Relative in vitro growth rates of duckweeds (Lemnaceae) – the most rapidly growing higher plants. Plant Biology 17: 33–41.

Züst, T., and A. A. Agrawal. 2016. Trade-Offs Between Plant Growth and Defense Against Insect Herbivory: An Emerging Mechanistic Synthesis. Annual Review of Plant Biology 68: 1–22.

