## Supplemental 1 for "Neopolyploidy-induced changes in the giant duckweed (*Spirodela polyrhiza*) alter herbivore preference, performance, and plant population performance"

**SUPPLEMENTARY MATERIAL**

Assour et al. – Appendix S1

**Table S1**: Genetic lineages of Spirodela polyrhiza used in this study and their corresponding latitude and longitude of the collection site.

| **Genetic Lineage** | **Sampling location** | **Latitude** | **Longitude** |
|---|---|---|---|
| SP.01 | Beaver County, PA, USA | 40.72981667 | -80.36023333 |
| SP.05 | Allegheny County, PA, USA | 40.621 | -79.82675 |
| SP.07 | Allegheny County, PA, USA | 40.62075 | -79.82813333 |
| SP.11 | Allegheny County, PA, USA | 40.62063 | -79.82718 |
| SP.41 | Butler County, PA, USA | 40.97156667 | -80.0186 |
| SP.43 | Andover Township, Ohio, USA | 41.606409 | -80.54012 |

**Table S2**: Surface area ($cm^2$) of single fronds from each of the six genetic lineages of *Spirodela polyrhiza* for diploids (2x) and neopolyploids (4x). Each estimate is a mean ± SE of 3 individuals.

| Genetic lineage | Ploidy | Mean surface area ($cm^2$) | Standard error ($cm^2$) |
|---|---|---|---|
| SP.01 | 2X | 0.52 | 0.035 |
| | 4X | 0.84 | 0.047 |
| SP.05 | 2X | 0.58 | 0.059 |
| | 4X | 1.12 | 0.14 |
| SP.07 | 2X | 0.97 | 0.11 |
| | 4X | 1.10 | 0.035 |
| SP.11 | 2X | 0.62 | 0.035 |
| | 4X | 0.97 | 0.065 |
| SP.41 | 2X | 0.53 | 0.044 |
| | 4X | 0.68 | 0.017 |
| SP.43 | 2X | 0.55 | 0.028 |
| | 4X | 0.79 | 0.072 |

**Table S3** Aphid abundances: ANOVA table showing the impact of ploidy (diploid, neopolyploid), duckweed genetic lineage and time-block on aphid abundance. Day and Jar ID (or experimental unit) were included as crossed random effects. Chi square values, df and P from a negative binomial generalized linear model are given.

| **Factor** | **Chisq** | **Df** | **Pr(>Chisq)** |
|---|---|---|---|
| Ploidy | 11.81 | 1 | 0.0031 |
| Lineage | 29.42 | 5 | <0.001 |
| Time-block | 41.36 | 1 | <0.001 |
| Ploidy:Lineage | 26.04 | 5 | <0.001 |

**Table S4** Lineage-specific negative binomial GLMMs for aphid abundances ANOVA table showing the impact of ploidy (diploid, neopolyploid) and time-block on aphid abundance. Day and Jar ID (or experimental unit) were included as crossed random effects. Chi square values, df and P from a negative binomial generalized linear model are given.

| Lineage | | Chisq | Df | Pr(>Chisq) |
|---|---|---|---|---|
| **SP.01** | Ploidy | 1.51 | 1 | 0.22 |
| | Time-block | 3.07 | 1 | 0.080 |
| **SP.05** | Ploidy | 0.81 | 1 | 0.29 |
| | Time-block | 1.08 | 1 | 0.23 |
| **SP.07** | Ploidy | 0.56 | 1 | 0.50 |
| | Time-block | 2.95 | 1 | 0.086 |
| **SP.11** | Ploidy | 4.13 | 1 | 0.049 |
| | Time-block | 3.60 | 1 | 0.15 |
| **SP.41** | Ploidy | 0.86 | 1 | 0.35 |
| | Time-block | 1.27 | 1 | 0.26 |
| **SP.43** | Ploidy | 0.0003 | 1 | 0.99 |
| | Time-block | 4.95 | 1 | 0.026 |

**Table S5** Duckweed abundances: ANOVA table showing the impact of plody (diploid, neopolyploid), duckweed genetic lineage, herbivory (present or absent) and time-block on duckweed abundance. Day and Jar ID (or experimental unit) were included as crossed random effects. Chi square values, df and P from a Poisson generalized linear model are given.

| **Factor** | **Chisq** | **Df** | **Pr(>Chisq)** |
|---|---|---|---|
| Ploidy | 73.67 | 1 | <0.001 |
| Lineage | 124.48 | 5 | <0.001 |
| Herbivory | 19.25 | 1 | <0.001 |
| Time-block | 20.13 | 1 | <0.001 |
| Ploidy:Lineage | 62.40 | 5 | <0.001 |
| Ploidy:Herbivory | 0.99 | 1 | 0.32 |
| Lineage:Herbivory | 24.63 | 5 | <0.001 |
| Ploidy:Lineage:Herbivory | 12.25 | 5 | 0.031 |

**Table S6** Lineage-specific Poisson GLMMs for duckweed abundances ANOVA table showing the impact of ploidy (diploid, neopolyploid), herbivory and time-block on duckweed abundance. Day and Jar ID (or experimental unit) were included as crossed random effects. Chi square values, df and P from a Poisson generalized linear model are given.

| Lineage | | Chisq | Df | Pr(>Chisq) |
|---|---|---|---|---|
| **SP.01** | Ploidy | 92.04 | 1 | <0.001 |
| | Herbivory | 25.07 | 1 | <0.001 |
| | Ploidy:Herbivory | 1.25 | 1 | 0.17 |
| | Time-block | 15.36 | 1 | <0.001 |
| **SP.05** | Ploidy | 99.04 | 1 | <0.001 |
| | Herbivory | 5.10 | 1 | 0.027 |
| | Ploidy:Herbivory | 10.38 | 1 | 0.0012 |
| | Time-block | 19.79 | 1 | <0.001 |
| **SP.07** | Ploidy | 1.96 | 1 | 0.18 |
| | Herbivory | 34.55 | 1 | <0.001 |
| | Ploidy:Herbivory | 0.38 | 1 | 0.56 |
| | Time-block | 0.075 | 1 | 0.77 |
| **SP.11** | Ploidy | 18.73 | 1 | <0.001 |
| | Herbivory | 8.63 | 1 | 0.0035 |
| | Ploidy:Herbivory | 0.45 | 1 | 0.56 |
| | Time-block | 8.44 | 1 | 0.002 |
| **SP.41** | Ploidy | 12.85 | 1 | <0.001 |

| | | | | |
|---|---|---|---|---|
| | Herbivory | 31.13 | 1 | <0.001 |
| | Ploidy:Herbivory | 0.96 | 1 | 0.52 |
| | Time-block | 1.14 | 1 | 0.31 |
| **SP.43** | Ploidy | 20.12 | 1 | <0.001 |
| | Herbivory | 33.12 | 1 | <0.001 |
| | Ploidy:Herbivory | 1.24 | 1 | 0.37 |
| | Time-block | 0.0064 | 1 | 0.95 |

**Table S7** Final duckweed biomass: ANOVA table showing the impact of ploidy (diploid or neopolyploid), herbivory (aphids or no aphids), genetic lineage and time-block on the final biomass of duckweed. Chi squared, df and P values given from the Gaussian generalized linear model.

| **Factor** | **Sum of Sq** | **Df** | **F value** | **Pr(>F)** |
|---|---|---|---|---|
| Ploidy | 0.00027 | 1 | 3.05 | 0.08 |
| Herbivory | 0.0020 | 1 | 23.3 | <0.001 |
| Lineage | 0.0020 | 5 | 4.55 | <0.001 |
| Time-block | 0.00079 | 1 | 9.10 | 0.0029 |
| Ploidy:Herbivory | 0.00030 | 1 | 3.49 | 0.063 |
| Ploidy:Lineage | 0.00042 | 5 | 0.97 | 0.43 |
| Lineage:Herbivory | 0.00032 | 5 | 0.74 | 0.59 |
| Ploidy:Lineage:Herbivory | 0.00092 | 5 | 2.11 | 0.065 |

**Table S8** Lineage-specific Gaussian linear models for final duckweed biomass ANOVA table showing the impact of ploidy (diploid, neopolyploid), herbivory and time-block on final duckweed biomass. Day and Jar ID (or experimental unit) were included as crossed random effects. Sum of square values, df, F-value and P-value from a linear model are given.

| Lineage | | Sum Sq | Df | F-value | Pr(>Chisq) |
|---|---|---|---|---|---|
| **SP.01** | Ploidy | 0.00027 | 1 | 1.88 | 0.18 |
| | Herbivory | 0.0020 | 1 | 14.39 | <0.001 |
| | Ploidy:Herbivory | <0.001 | 1 | 2.15 | 0.15 |
| | Time-block | <0.001 | 1 | 0.62 | 0.44 |
| **SP.05** | Ploidy | <0.001 | 1 | 1.11 | 0.30 |
| | Herbivory | 0.0014 | 1 | 17.67 | <0.001 |
| | Ploidy:Herbivory | <0.001 | 1 | 2.99 | 0.093 |
| | Time-block | <0.001 | 1 | 6.94 | 0.012 |
| **SP.07** | Ploidy | <0.001 | 1 | 1.32 | 0.26 |
| | Herbivory | 0.0022 | 1 | 31.35 | <0.001 |
| | Ploidy:Herbivory | <0.001 | 1 | 0.027 | 0.89 |
| | Time-block | <0.001 | 1 | 5.26 | 0.028 |
| **SP.11** | Ploidy | <0.001 | 1 | 0.026 | 0.87 |
| | Herbivory | <0.001 | 1 | 5.15 | 0.030 |
| | Ploidy:Herbivory | <0.001 | 1 | 2.76 | 0.11 |
| | Time-block | <0.001 | 1 | 7.50 | 0.010 |
| **SP.41** | Ploidy | <0.001 | 1 | 0.50 | 0.82 |

| | | | | | |
|---|---|---|---|---|---|
| | Herbivory | 0.0012 | 1 | 37.40 | <0.001 |
| | Ploidy:Herbivory | <0.001 | 1 | 0.19 | 0.66 |
| | Time-block | <0.001 | 1 | 1.10 | 0.30 |
| **SP.43** | Ploidy | 0.0010 | 1 | 0.24 | 0.63 |
| | Herbivory | <0.001 | 1 | 33.41 | <0.001 |
| | Ploidy:Herbivory | <0.001 | 1 | 0.92 | 0.35 |
| | Time-block | <0.001 | 1 | 0.050 | 0.83 |
